# Loquacious Modulates Flaviviral RNA Replication in Mosquito Cells

**DOI:** 10.1101/2021.12.03.471060

**Authors:** Shwetha Shivaprasad, Kuo-Feng Weng, Yaw Shin Ooi, Julia Belk, Jan E. Carette, Ryan Flynn, Peter Sarnow

## Abstract

Arthropod-borne viruses infect both mosquito and mammalian hosts. While much is known about virus-host interactions that modulate viral gene expression in their mammalian host, much less is known about the interactions that involve inhibition, subversion or avoidance strategies in the mosquito host. A novel RNA-Protein interaction detection assay was used to detect proteins that directly or indirectly bind to dengue viral genomes in infected mosquito cells. Membrane-associated mosquito proteins SEC61A1 and Loquacious (Loqs) were found to be in complex with the viral RNA. Depletion analysis demonstrated that both SEC61A1 and Loqs have pro-viral functions in the dengue viral infectious cycle. Co-localization and pull-down assays showed that Loqs interacts with viral protein NS3 and both full-length and subgenomic viral RNAs. While Loqs coats the entire positive-stranded viral RNA, it binds selectively to the 3’ end of the negative-strand of the viral genome. In-depth analyses showed that the absence of Loqs did not affect translation or turnover of the viral RNA but modulated viral replication. Loqs also displayed pro-viral functions for several flaviviruses in infected mosquito cells, suggesting a conserved role for Loqs in flavivirus-infected mosquito cells.

**Author Summary:** There is a wealth of information that dictates virus-host interactions in flavivirus-infected mammalian cells, yet there is only sparse information on the mechanisms that modulate viral gene expression in the mosquito host. Using a novel RNA-protein detection assay, the interactions of SEC61A1 and Loqs with the dengue viral genome were found to have proviral functions in infected mosquito cells. In particular, Loqs forms complexes with the positive-strand of the viral RNA and the very 3’ end of the negative-strand viral RNA. Further analyses showed that Loqs modulates viral RNA replication of dengue virus and gene amplification of several other flaviviral genomes. These findings argue that Loqs is an essential proviral host factor in mosquitos.

## Introduction

Dengue virus (DENV) is an enveloped, single-stranded positive sense RNA virus belonging to the *Flaviviridae* family. It infects ∼400 million people worldwide every year and is transmitted by the *Aedes aegypti* and *Aedes albopictus* species of mosquitoes [1]. The ∼11kb DENV genomic RNA consists of an open reading frame that codes for the structural and nonstructural viral proteins, flanked by 5’ and 3’ untranslated regions (UTR) (Fig. 1A). The viral UTRs are involved in multiple RNA-RNA and RNA-protein interactions that regulate the efficiency of infection, modulate host innate immune responses and viral pathogenesis [2, 3].

**Fig 1.**
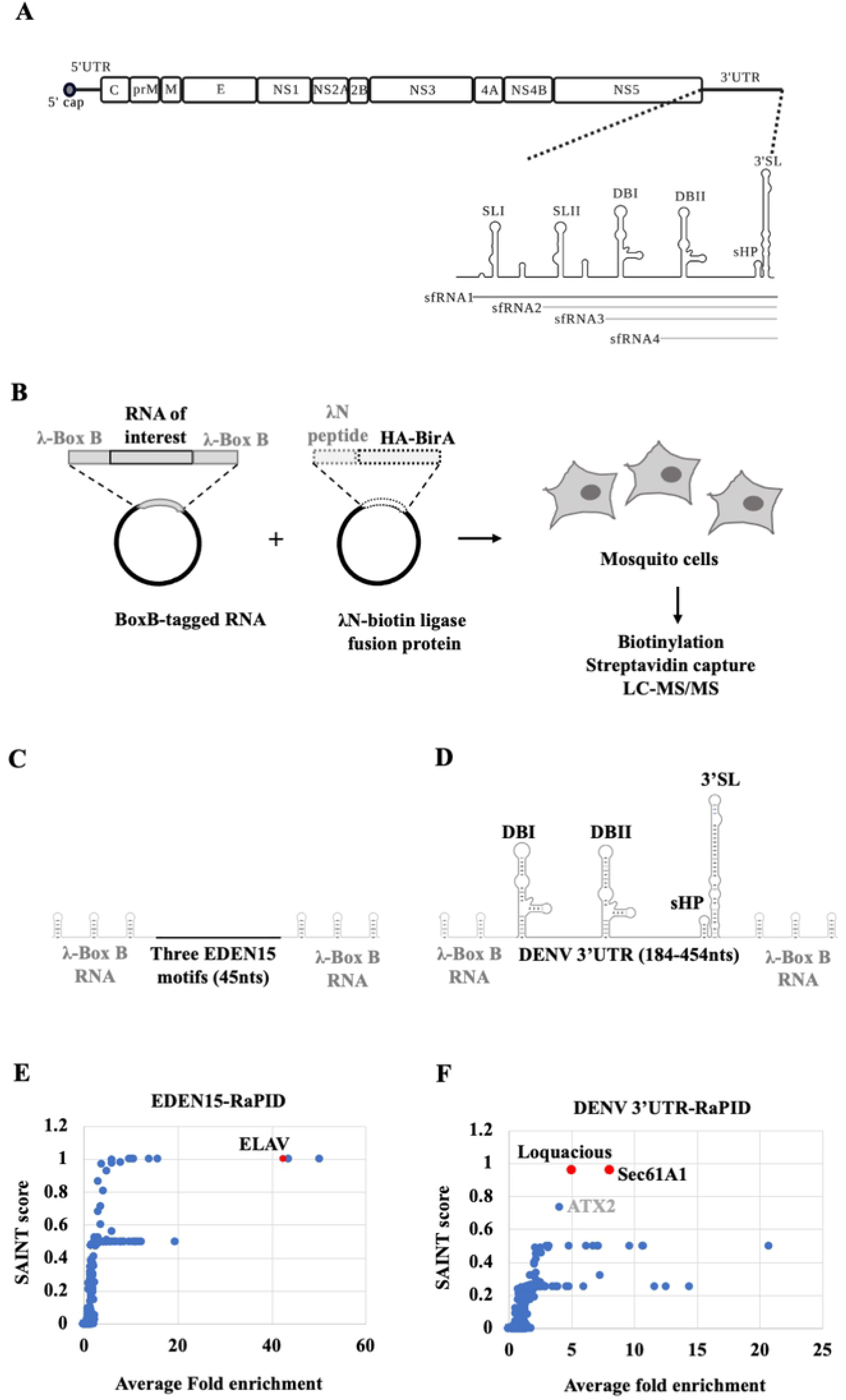
Diagram of the dengue viral (DENV) genome and strategy for the RNA-protein interaction (RAPID) assay. (A) DENV genome organization. The open reading frame encoding the three structural (C (capsid), prM/M (membrane), E (envelope)) proteins and seven non-structural (NS1, NS2A, NS2B, NS3, NS4A, NS4B, NS5) proteins flanked by 5’ and 3’ untranslated regions (UTR) are shown. The 3’ UTR is organized into stem loops SLI, SLII, dumbbell structures DBI, DBII, a small hairpin (sHP) and a terminal 3’ stem loop (3’SL). Subgenomic RNA fragments (sfRNA) 1-4 are indicated. (B) Outline of RaPID assay. Plasmids expressing BoxB-flanked DENV2 of sfRNA3/4 and the λN-biotin ligase fusion protein gene (λN-HA-BirA) were co-transfected into mosquito cells. Subsequently, biotinylated proteins were captured using streptavidin beads and identified by LC-MS/MS. (C) Schematic of the EDEN15 RNA motifs (3 repeats of 15bp each) flanked by three BoxB RNA motifs each at their 5’ and 3’ ends. (D) DENV2 3’UTR (184-454nts) flanked by two BoxB RNA motifs at its 5’ end and three BoxB motifs at its 3’ end. (E) Average fold change of proteins enriched in EDEN15 RNA expressing cells relative to the scrambled RNA control plotted against their SAINT probability scores. ELAV protein (shown in red) was enriched by ∼40 fold (n=2, **p<0.005). (F) Average fold change of proteins enriched in DENV 3’UTR expressing cells relative to the scrambled RNA control is plotted against their SAINT probability scores. Sec61A1 and Loquacious proteins (shown in red) were enriched by ∼8 fold and ∼5 fold respectively (n=4, *p<0.05, **p<0.005).

The interactions of the DENV 3’UTR with viral and host cellular proteins are particularly interesting because of the diverse roles of the 3’UTR in viral infection. First, it is the site of initiation of viral RNA replication. Secondly, it is a hotspot for accumulation of adaptive mutations in both human and mosquito hosts. Third, it is resistant to degradation by host exoribonuclease XRN1, which allows subgenomic flaviviral RNAs (sfRNAs) to accumulate (Fig 1A) [4–6]. Biochemical methods such as RNA-affinity capture have identified several human proteins in complexes with the 3’ end of DENV genomic RNA or sfRNAs to regulate viral replication and immune evasion [7–11]. A recent study used ChIRP-MS (Comprehensive Identification of RNA Binding Proteins by Mass spectrometry), an intracellular crosslinking approach to identify numerous RNA binding proteins that interact directly with the DENV RNA in mammalian cells [12]. Specifically, the endoplasmic reticulum-associated proteins vigilin and ribosome-binding protein 1 (RRBP1) were associated with DENV RNA, and modulated viral RNA replication and translation, respectively [8].

While most studies have focused on viral RNA-host protein interactions in mammalian cells, there has been less investigation of proteins that form complexes with the DENV RNA and regulate infection in mosquito cells [3]. In this study, we have employed an intracellular biotinylation-based approach called RaPID (RNA-Protein Interaction Detection)[13] to identify mosquito proteins that form complexes with the DENV 3’UTR in mosquito cells. While this technique has been previously used to identify proteins that bind to Zika virus and rotavirus in mammalian cells [14, 15], adaptation to mosquito cells has allowed us to identify critical regulators of DENV replication in the mosquito host.

## Results

### Biotinylation-based proteome analysis identifies DENV 3’UTR-protein interactions in mosquito cells

We designed the RaPID system to identify proteins that interact with the dengue viral (New Guinea strain DENV2-NGC, DENV) 3’UTR (Fig. 1A,B) in mosquito cells. A 271 nucleotide (nt) region at the 3’ end of the DENV 3’UTR (184-454 nts) was chosen as a target as it is present both in the genomic RNA as well as the subgenomic RNA fragments, sfRNA3 and sfRNA4, which specifically accumulate in mosquito cells during infection with mosquito-adapted DENV RNA, whereas sfRNAs1/2 predominately accumulate in infected human cells [6, 16]. Briefly, the target DENV RNA sequence, encompassing sfRNAs 3/4 (Fig. 1A), was expressed within flanking phage λN-BoxB RNA motifs, together with a fusion protein composed of biotin ligase and the λN-BoxB RNA binding protein in mosquito cells (Fig. 1B). In the presence of biotin, the λN-biotin ligase is expected to bind with the BoxB RNA stem loops and biotinylate proteins that bind directly or are in a complex with the target DENV RNA sequence. Biotinylated proteins can then be isolated using streptavidin beads and identified by mass spectrometry (LC-MS/MS) (Fig. 1B).

In a proof-of-principle experiment, we tested whether RaPID could detect the known RNA-protein complex between the EDEN15 RNA motif and ELAV family of proteins [15] in mosquito C6/36 cells. C6/36 cells were co-transfected with plasmids expressing the EDEN15-BoxB RNA (Fig. 1C) and the λN-biotin ligase protein. As a negative control, cells were co-transfected with plasmids expressing a scrambled EDEN15 sequence and the λN-biotin ligase protein. Biotinylated proteins were isolated using streptavidin beads and the peptides were identified by LC-MS/MS. Results were filtered and peptides that were enriched at the EDEN15 motif with a probability score (SAINT score) of greater than 0.9 relative to the scrambled sequence were shortlisted as true binders (**Table S1**). We identified the mosquito ELAV protein, a homolog of human CELF1 to be ∼40-times enriched in the EDEN15 expressing cells relative to the scrambled sequence, suggesting an interaction of the ELAV protein with the EDEN15 RNA (Fig. 1E). This experiment confirmed that the RaPID pipeline works efficiently in mosquito cells.

Next, we expressed the DENV 3’ UTR-BoxB RNA (Fig 1D) and the λN-biotin ligase protein in mosquito cells. RaPID analysis (**Table S1**) identified two high confidence hits (Fig.1F) that could potentially interact with this region of the DENV 3’UTR in mosquito cells: AAEL010716 is an unannotated gene in mosquitoes, but is 90% identical to the *Drosophila* endoplasmic reticulum transport protein Sec61 subunit alpha (Sec61A1) [17]. AAEL008687 (*Loquacious, Loqs*) encodes a dsRNA-binding protein Loqs. Curiously the Loqs-PA isoform is involved in the RNAi immune response pathway in mosquitos [18]. Sec61A1 is known to play a proviral role in flaviviral infection in both human and mosquito cells by modulating viral mRNA translation [19]. However, no role for Loqs in viral infection has been suspected. In addition, RaPID identified A0A182HDU4 encoding ATX2, which displays 24% identity to human ATX2. ATX2 has been shown to bind to several DEAD box RNA helicases, which are known to be involved in RNA processing pathways both in mosquitos and humans [20, 21]. However, ATX2 was enriched with a slightly lower SAINT probability score of 0.736 (Fig. 1F). We chose to pursue the potentially novel roles for Sec61A1 and Loqs in DENV-infected mosquito cells.

### Partial depletion of Sec61A or Loqs inhibits DENV replication in mosquito cells

To determine the effects of Sec61A1 and Loqs on DENV RNA expression, mRNAs encoding these proteins were depleted in infected cells. Because C6/36 cells are deficient in the RNAi pathway [22], RNAi-competent Aag2 mosquito cells [23] were used in the mRNA depletion assays. Loqs has two abundant isoforms in mosquito cells, Loqs-PA and Loqs-PB. Thus, dsRNAs were designed to target both isoforms (dsLoqs) or only the Loqs-PB isoform (dsLoqs-PB), which can be selectively targeted due to the presence of a unique exon 5. Aag2 cells were transfected with 500bp long double-stranded RNAs targeting Sec61A1 or Loqs mRNAs. Transfected cells were subsequently infected with the DENV2-New Guinea C strain (NGC) at a multiplicity of infection (MOI) of 0.1. At 96 hrs post infection, cells were harvested and processed for downstream assays (Fig. 2A).

**Fig 2.**
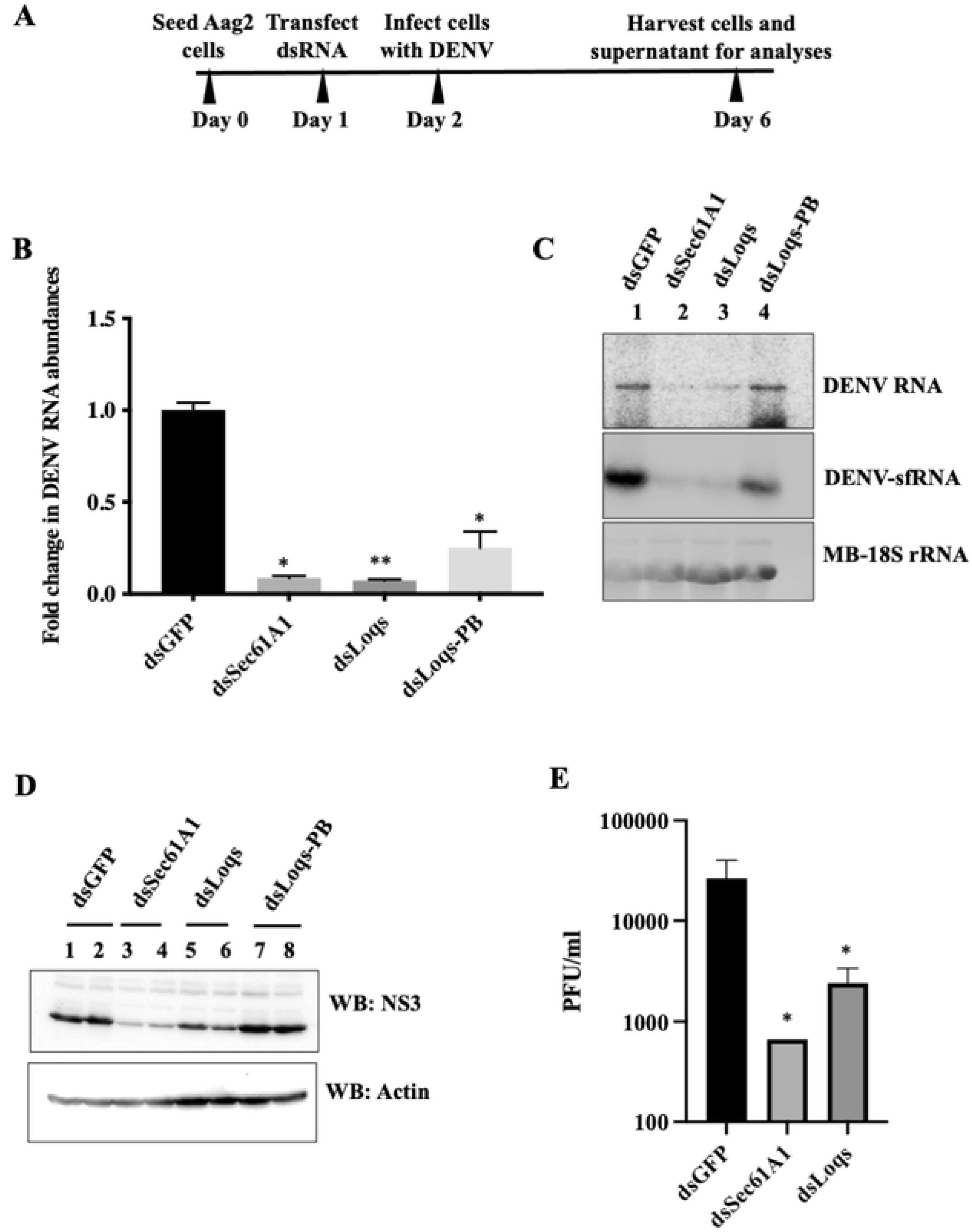
Effects of Sec61A1 and Loquacious depletion on DENV2 RNA and protein abundances, and viral titers. (A) Experimental outline. Mosquito Aag2 cells were transfected with double stranded RNAs (dsRNA) directed against GFP, Sec61A1, Loqs (targeting both PA and PB isoforms) or Loqs-PB mRNAs. 24 hrs post transfection, cells were infected with DENV2-NGC at an MOI of 0.1 and harvested 96 hrs post infection for analyses. (B) RT-qPCR measurement of DENV RNA abundances in dsRNA-treated cells plotted as fold change over treatment with dsGFP. Data was normalized to internal control RPL32 mRNA levels (n=3, *p<0.05, **p<0.005). (C) Effects of dsRNA treatment on DENV RNA and DENV subgenomic RNA (DENV-sfRNA) abundances, measured by Northern blot analysis. Methylene blue (MB) staining of rRNA was used as a loading control. Representative image from three independent experiments is shown. (D) Effects of dsRNA treatment on DENV NS3 and actin protein levels examined by western blot analysis. Representative image from three independent experiments is shown. (E) Effects of dsRNA treatment on infectious viral titers determined by plaque forming assays (PFU/ml, n=4, *p<0.05).

Efficiencies of Sec61A1 and Loqs mRNA depletions were demonstrated by measuring the individual mRNA abundances by qPCR (Fig. S1A). Depletion of specific isoform proteins of Loqs was also validated by western blot analysis (Fig. S1B). RT-PCR analysis revealed a significant reduction in DENV RNA abundances upon depletion of either Sec61A1 or Loqs mRNAs (Fig. 2B). Depleting both isoforms of Loqs (dsLoqs) had a greater effect on viral RNA levels than depleting just the PB isoform (dsLoqs-PB), suggesting that Loqs-PA might play a more predominant role in supporting viral infection (Fig. 2B). Northern blot analysis using probes against the DENV 3’UTR indicated a reduction of both DENV genomic RNA and sfRNA abundances in Sec61A1 and Loqs depleted cells relative to the control dsGFP-treated cells (Fig. 2C). Depletion of Sec61A1/Loqs also resulted in a significant reduction in DENV NS3 protein abundance (Fig. 2D). Finally, plaque assays indicated a significant hundred-fold reduction in the viral titer in both dsSec61A1-and dsLoqs-treated cells (Fig. 2E). These data argue that both Sec61A1 and Loqs have pro-viral functions in DENV-infected mosquito cells.

To examine whether the observed effects were specific to DENV2-NGC, mRNA depletion experiments were repeated in cells infected with Thailand strain DENV2-16681, which has distinct mutations from the DENV2-NGC virus [24]. Results in Fig. S1C showed that depletion of SEC61A1 or Loqs also diminished DENV2-16681 RNA abundances. Similarly, a reduction in both extracellular (Fig. S1D) and intracellular (Fig. S1E) viral RNA abundances was observed after depletion of SEC61A1 or Loqs, arguing that that both proteins have pro-viral functions in the infectious cycle in both DENV2-NGC and DENV2-16681 infected cells.

In order to rule out any non-specific effects associated with using dsRNAs, which are processed into multiple siRNAs, we designed individual siRNAs targeting either Sec61A1 or Loqs mRNAs. A significant reduction in DENV RNA levels in Sec61A1- and Loqs-siRNA-treated cells was observed relative to the non-targeting control siRNA treated cells (Fig S2A). Because it is unlikely that all siRNAs have the same off-target effect, this data argues that dsRNAs specifically depleted SEC61A1 and Loqs mRNAs. To rescue the phenotype observed after depletion of SEC61A1 and Loqs, we expressed Loqs-PA and Loqs-PB encoding cDNAs in cells treated with siRNAs targeting the 3’UTR of endogenous Loqs. However, no rescue of the siLoqs-induced phenotype was observed (Fig. S2B). Unfortunately, we were also unable to express the full length Sec61A1 protein in our rescue studies.

To test if these host proteins are essential in the infectious cycles of other RNA viruses, Sec61A1 and Loqs-depleted Aag2 cells were infected with West Nile virus (WNV), yellow fever virus (YFV), Zika virus (ZIKV) or Chikungunya virus (CHIKV). A significant reduction in YFV and ZIKV RNA abundances were observed after depletion of either of these two proteins (Fig. 3). In case of WNV, depletion of Sec61A1 affected viral RNA abundances, while depletion of Loqs had no effect (Fig. 3). Surprisingly, depletion of either of the two proteins resulted in an increase in CHIKV RNA abundances, a virus which belongs to the *Togaviridae* family. These experiments suggest that Sec61A1 and Loqs play a proviral role in the infectious cycle of several flaviviruses, but can play antiviral roles as well, as is often seen in the battle between viruses and their hosts. Next, we focused on characterizing the role of Loqs in viral infection in more detail.

**Fig 3.**
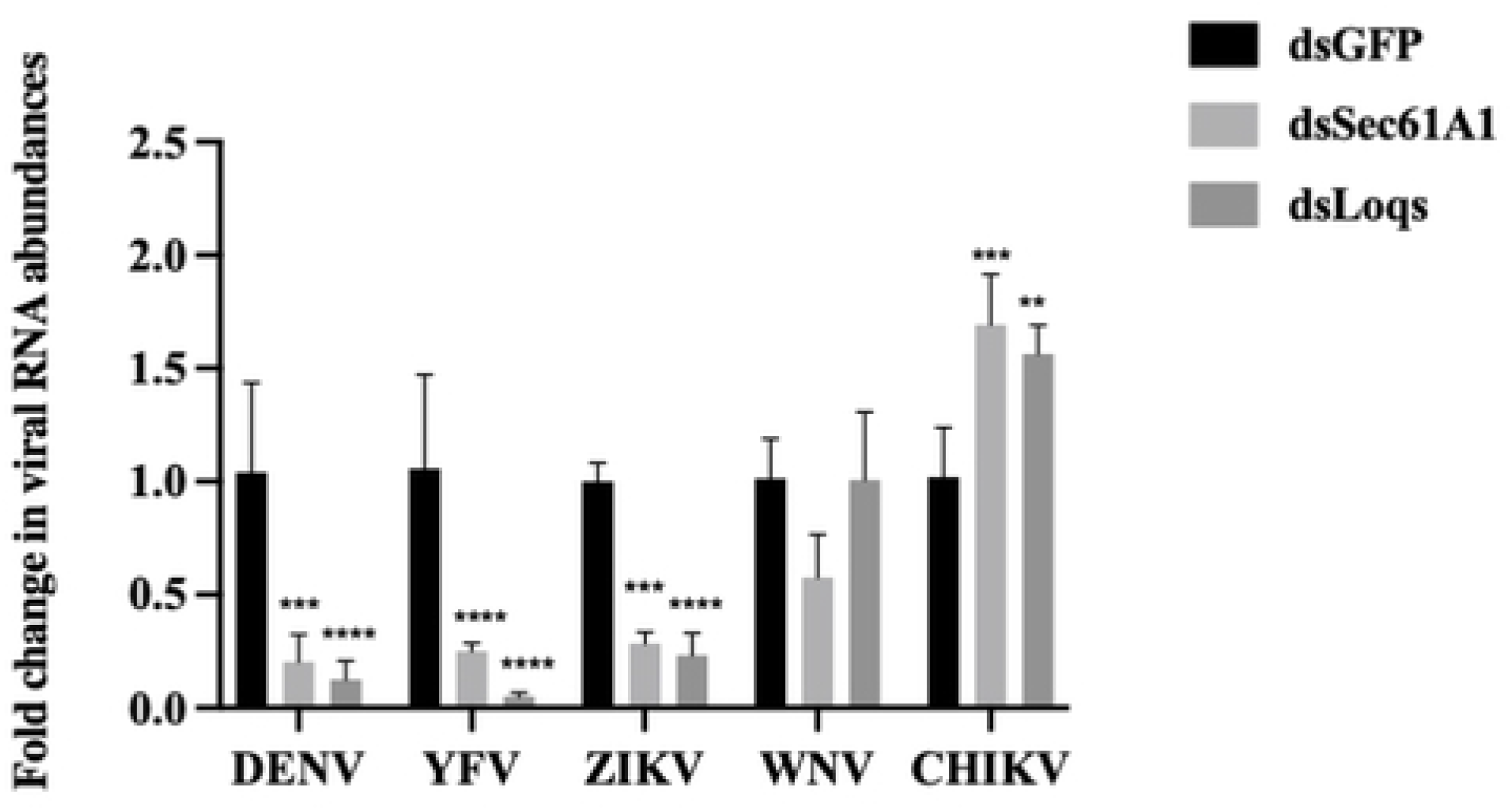
Effects of Loqs depletion distinct RNA virus infections. Aag2 cells were infected with dengue virus (DENV), West Nile virus (WNV), yellow fever virus (YFV), Zika virus (ZIKV) or chikungunya virus (CHIKV) at a MOI of 0.1 and harvested at 96 hrs post infection. Viral RNA abundances were measured by qPCR using specific primers. Data is represented as average fold-change over dsGFP from three independent experiments.

### Loqs colocalizes and forms complexes with DENV RNA in infected mosquito cells

The RaPID approach suggested that Loqs directly or indirectly binds to the DENV genomic/subgenomic RNA sequences. To test if Loqs co-localizes with the genomic DENV RNA in infected cells, in situ hybridization experiments were carried out in DENV-infected Aag2 cells. Immunostaining for endogenous Loqs protein showed that it is predominantly located in the cytoplasm in infected cells (Fig. 4A). DENV RNA was visualized by in situ RNA hybridization, using fluorescently labeled probes directed against viral NS5 region, which also localized to the cytoplasm. To examine whether Loqs and DENV RNA significantly colocalized, images were analyzed by Color 2, an algorithm for measuring colocalization in pixel images, and the significance of colocalizations was determined by the Costes P-value [25]. The results showed that Loqs protein and DENV RNA significantly colocalized with each other (Costes P-value > 0.95) in 80% (28/35) of the analyzed infected (DENV RNA positive) cells (Fig 4A**, lower panels)**. In comparison, the known colocalization of DENV NS3 protein with DENV RNAs [26] was 97% (34/35) of the inspected cells. These results suggest that Loqs colocalizes with DENV RNAs with a significance that is comparable to that of DENV RNA-NS3 colocalization.

**Fig 4:**
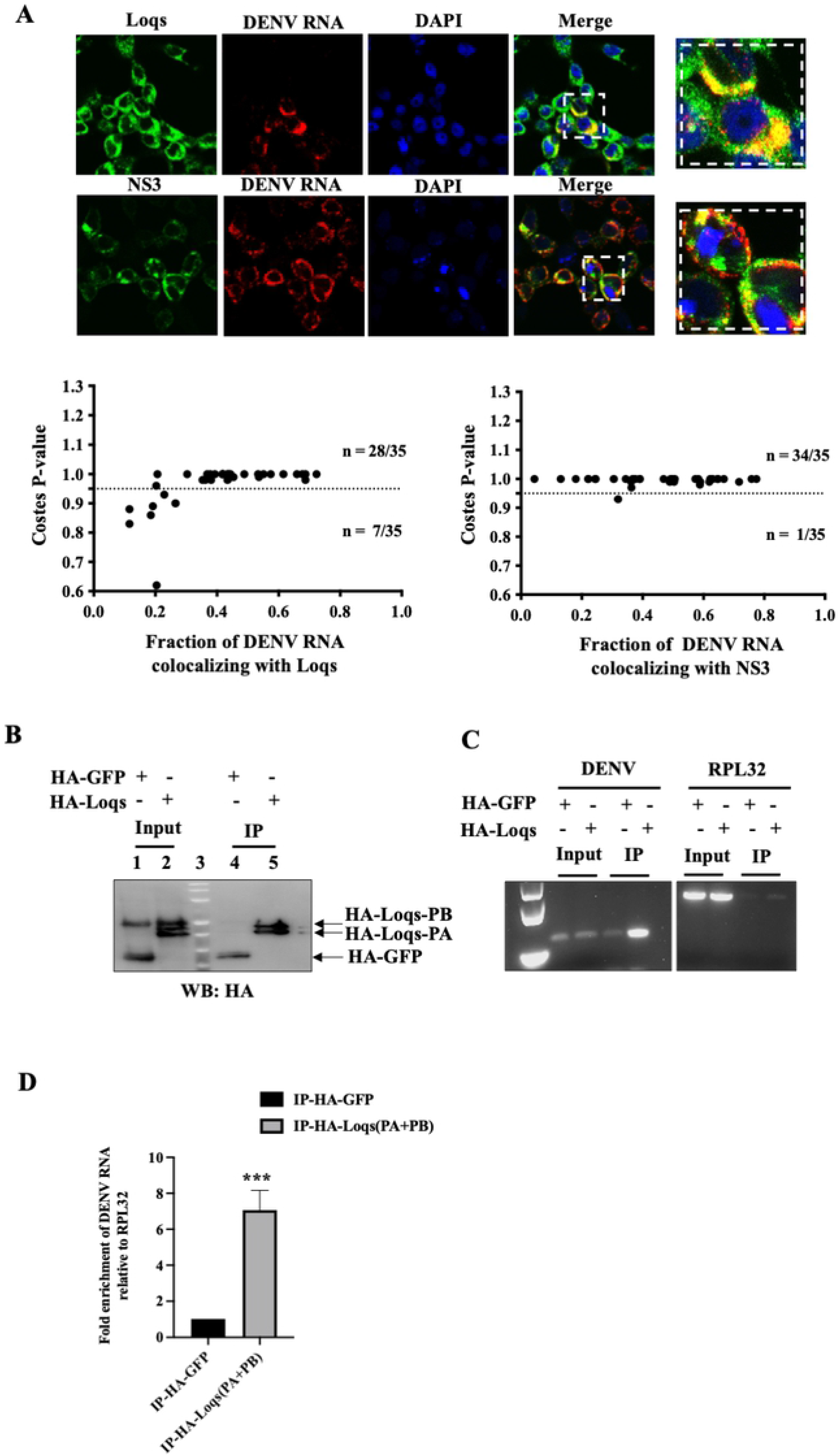
Colocalization and interaction of Loqs protein with DENV RNA. (A) Fluorescent in situ hybridization imaging of Aag2 cells infected with DENV2 at an MOI of 1 after 48 hrs. NS3 and Loqs proteins (shown in green) were visualized using labeled antibodies, while DENV RNA (shown in red) was visualized using labeled antisense RNA probes. Costes p value was calculated to measure the extent of colocalization of DENV2 RNA with NS3/Loqs proteins. (B) Immunoprecipitation of HA-tagged Loqs from Aag2 cells infected with DENV2 at a MOI of 1. Aag2 cells transfected with HA-GFP or HA-Loqs PA/PB plasmids were infected with DENV2 for 48h, and immunoprecipitations were performed with anti-HA antibodies. Abundances of HA-GFP and HA-Loqs in input lysates and immunoprecipitated material measured by western blot analysis. (C) DENV2 and RPL32 RNA abundances in immunoprecipitated RNA (IP) and input RNA (10%) were measured by semi-quantitative RT-PCR. A representative agarose gel image from three independent experiments is shown. (D) DENV2 and Loqs mRNA abundances in immunoprecipitated RNA (IP) and input RNA (10%) measured by RT-qPCR. Data was normalized to RPL32 mRNA levels (n=3, ****p=0.0006).

To detect Loqs protein-DENV RNA complexes by immunoprecipitation, Aag2 cells were transfected with HA-tagged Loqs PA or HA-EGFP and subsequently infected with DENV infection. HA-tagged proteins were immunoprecipitated (Fig. 4B) and DENV RNA was detected by semi-quantitative PCR (Fig 4C) and qPCR (Fig 4D). A significant enrichment of DENV RNA was observed upon HA-Loqs immunoprecipitation as compared to the control immunoprecipitations. Northern blot analysis showed that Loqs binds both to the full-length DENV RNA and to sfRNAs, relative to the IgG control (Fig. S3A). To pinpoint the region on the viral RNA where Loqs binds, we performed infrared crosslinking and immunoprecipitation (irCLIP) assays [13] in infected cells expressing HA-Loqs. Reverse transcriptase stops for Loqs were mapped across the full-length viral RNA including the UTRs (Fig. S3B). The results showed that Loqs interacted with the entire positive-stranded viral RNA, with a few dozen hot spots. However, Loqs poorly interacted with the negative-stranded viral RNA with an exceptional single specific band at the very 3’ end of the negative strand (Fig. S3C). These findings show that Loqs can coat the entire positive-strand viral RNA, possibly by its relative accessibility.

### Loqs modulates DENV RNA replication

Because Loqs supports viral infection and interacts with DENV RNA, we tested whether Loqs affects translation, replication or stability of the viral RNA. First, we tested if Loqs is associated with the endoplasmic reticulum (ER), which is the primary site for DENV RNA translation and replication. A digitonin-based fractionation method was employed to separate the cytoplasmic and membrane-associated proteins and to determine the localization of Loqs by western blot analysis. Both Loqs-PA and Loqs-PB were enriched in the membrane fractions in both uninfected and infected cells (Fig. 5A), as was the DENV NS3 protein in infected cells. Next, the association of Loqs with replication proteins NS3, NS5, NS4B and the viral capsid proteins were studied. Immunoprecipitation of HA-tagged Loqs-PA from infected Aag2 cells indicated complex formation with only NS3 protein (Fig. 5B), which is essential for both viral RNA translation and replication [26]. This complex formed with or without RNAse treatment (Fig. 5B). These findings argue that Loqs is specifically associated with NS3 in membranes in infected cells.

**Fig 5.**
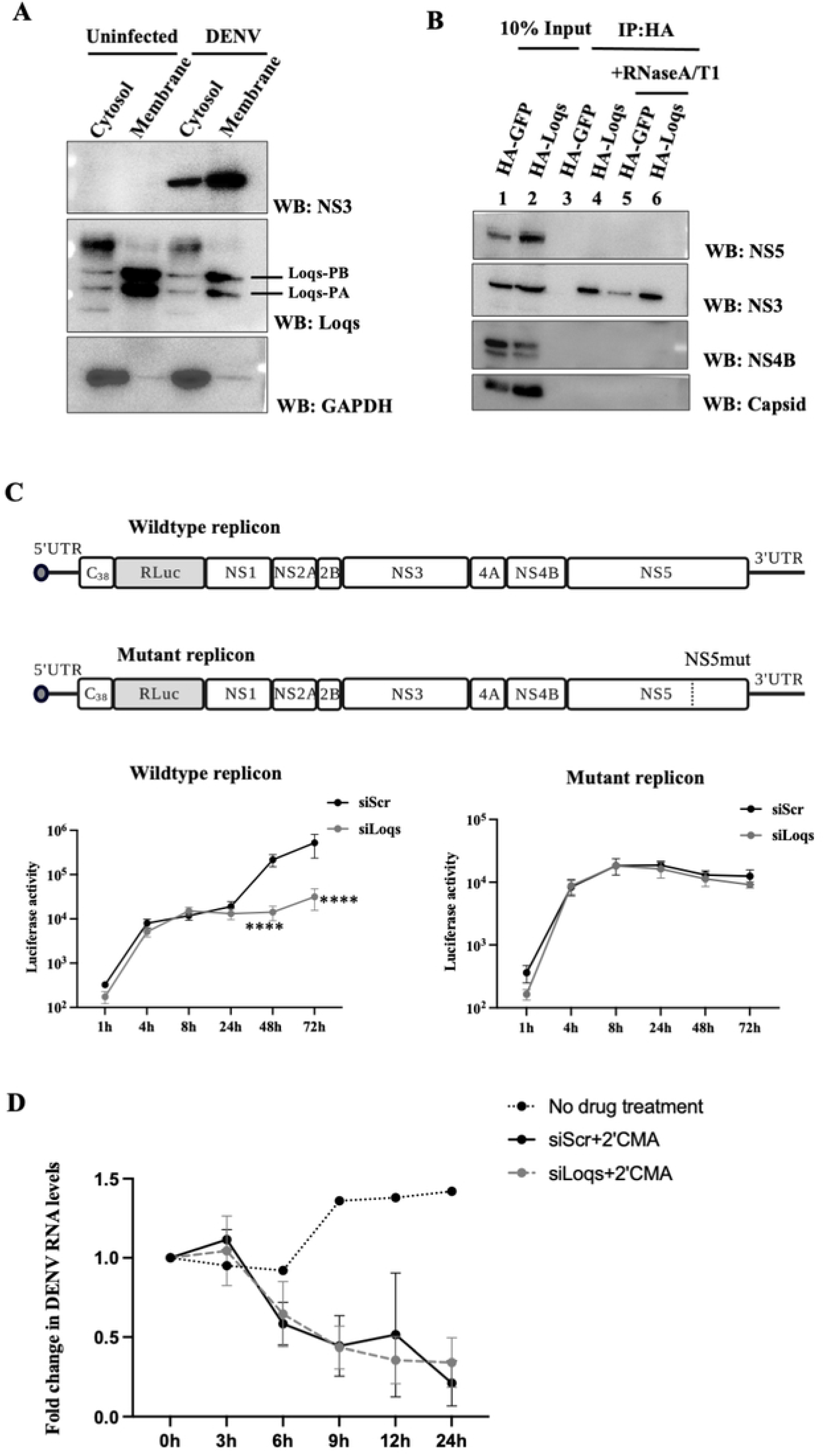
Effect of Loqs depletion on DENV RNA translation, replication and stability. (A) Western blot analysis of NS3, Loqs and GAPDH protein abundances in cytosolic and ER membrane fractions isolated from DENV2-infected Aag2 cell lysates at 72 hrs post infection. Representative image from three independent experiments is shown. (B) Immunoprecipitation of HA-tagged Loqs from Aag2 cells infected with DENV2 at a MOI of 1. Aag2 cells transfected with HA-GFP or HA-Loqs PA/PB plasmids were infected with DENV2 for 72 hrs and immunoprecipitations were performed with anti-HA antibody with or without RNaseA/T1 treatment. The abundances of DENV NS3, NS4B and capsid proteins in the immunoprecipitated material (IP) and the input lysates (10%) were determined by western blot analysis. (C) Luciferase activities of wildtype and replication-defective DENV2 luciferase replicons in C6/36 cells transfected with control siRNAs or siRNAs against Loqs (siLoqs-4 and siLoqs-5 were used at a final concentration of 25nM each). C6/36 cells were transfected with the indicated siRNAs followed by wildtype or replication-defective (NS5-GDD) replicon RNAs and harvested at the indicated time points. Average luciferase expression from DENV replicons from six independent replicates is shown. (D) Effect of Loqs depletion on DENV RNA stability. Aag2 cells were transfected with the indicated siRNAs and infected with DENV2-NGC at MOI of 1. 24 hrs post infection, cells were treated with 20 μM 2’CMA to inhibit viral replication. Viral RNA abundances at different times post 2’CMA treatment were measured by qPCR. Data is represented as an average of two independent experiments.

To test if Loqs depletion affects viral RNA translation, the association of DENV RNA with polysomes in infected cells was examined. The abundance of DENV RNA in each individual fraction from cells treated with dsGFP or dsLoqs RNAs, was analyzed by qPCR (Fig. S4). DENV RNA was distributed in fractions 8 through 14 in both wildtype and Loqs-depleted cells suggesting that Loqs doesn’t affect the association of ribosomes with viral RNA, and the association of multiple ribosomes with individual RNAs (Fig. S4) argues that translation elongation is also not blocked when Loqs is depleted.

To examine effects of Loqs on viral RNA translation and replication in more detail, expression of luciferase-containing wildtype and replication-defective replicon RNAs were examined in Loqs-depleted C6/36 cells. C6/36 cells were used in this experiment as they were able to better support replicon expression as compared to Aag2 cells. The accumulation of viral RNA is affected by both its synthesis and degradation. We observed a one-log reduction in luciferase expression from the wildtype replicon in Loqs siRNA-transfected cells (Fig. 5C). However, there was no difference in luciferase expression from the replication-defective mutant in Loqs siRNA-treated cells (Fig. 5C), suggesting that Loqs primarily affects viral RNA replication rather than RNA translation. Furthermore, this result also confirms that the Loqs siRNAs did not display off-target effects on the viral genome.

The rate of degradation of DENV RNA in siLoqs-treated cells was examined after addition of the NS5 RNA polymerase inhibitor 2′-C-methyladenosine (2’CMA) [27, 28]. C6/36 cells were transfected with siRNAs, infected with DENV and viral RNA abundances were measured by qPCR. There was no significant difference in the rate of degradation of viral RNA in Loqs siRNA-treated DENV2 infected cells compared to the control siRNA-treated cells indicating that Loqs doesn’t affect viral RNA degradation (Fig. 5D). These experiments point towards a role for Loqs in DENV RNA replication, but not viral RNA stability.

### Effects of Loqs on DENV replication is independent of its role in the RNAi pathway

It is known that Loqs interacts with Dicer and Argonaute proteins to regulate both siRNA and miRNA pathways in mosquito cells [18]. Thus, we investigated whether Loqs could bind to the viral RNA and possibly protect the viral genome from siRNA or miRNA mediated degradation. To test this hypothesis, Dicer-2 KO Aag2 cells (AF319) cells [29] were treated with siRNAs directed against Sec61A1 or Loqs and transfected with infectious DENV-luciferase virus (Fig. S5A) or DENV-luciferase replicon RNAs (Fig. S5B). Depletion of Loqs in AF319 cells also resulted in a significant reduction of luciferase expression from DENV full length or replicon RNAs, suggesting that Loqs regulation of DENV replication is independent of Dicer-2.

## Discussion

Interactions of RNA viral genomes with host RNA binding proteins are essential for viral infection and immune evasion in both human and mosquito hosts. Host RNA-binding proteins interactions with viral RNA can result in structural and/or functional changes that can be either restrictive or supportive of viral infection. A recent ChIRP-MS screen identified RRBP and vigilin as DENV RNA binding proteins that support viral RNA translation, replication and stability [12]. In addition, the DENV 3’UTR is known to interact with and co-opt DDX6 and Lsm1 proteins to support viral RNA replication while other 3’UTR interacting proteins such as Quaking (QKI) play antiviral roles and inhibit DENV RNA replication [8, 9].

In addition to the DENV genomic RNA, DENV sfRNAs derived from the 3’UTR can also form complexes with host proteins to modulate viral transmission and immune evasion. For example, sequestration of TRIM25, G3BP and Caprin proteins by DENV sfRNAs suppresses antiviral interferon responses in mammalian cells while sfRNA interactions with Dicer and Ago2 proteins suppresses antiviral RNAi response in both mammalian and mosquito cells [10, 11]. A recent study showed that sequestration of the mosquito antiviral proteins ME31B, ATX2 and AAEL018126 by ZIKV and WNV sfRNAs increases viral transmission in mosquitoes [30].

Furthermore, viruses can target host RNA-binding proteins to modulate viral gene amplification. Our study identified Sec61A1 and Loquacious as proteins that interact with the DENV 3’UTR in mosquito cells. While several proteins associated with the ER have been shown to be critically important for viral replication in mammalian cells, the exact mechanism by which they regulate viral infection is unknown. Sec61A and Sec61B have been predicted to have non-canonical RNA binding activity, which could help in transporting the viral RNA to the ER for translation/replication [31, 32]. The identification of mosquito Sec61A1 in our RAPID screen suggests a possible direct interaction between Sec61A1 and the viral RNA which could influence viral RNA localization and/or replication.

We observed that Loqs colocalizes with NS3 in membranous fractions in DENV-infected mosquito cells and interacts with both DENV genomic and subgenomic RNAs. The lack of obvious hydrophobic or transmembrane regions in the protein suggests that Loqs could be anchored to the membranes by other membrane-associated host or viral proteins. The membrane localization of Loqs puts it at a strategic position to support viral translation and replication which occur in ER-derived membranous scaffolds. Because we did not observe any obvious effects of Loqs depletion on polysome association of the DENV RNA or luciferase expression from a non-replicating DENV replicon, we conclude that Loqs predominantly affects viral RNA replication. Because the stability of DENV RNA was not affected in Loqs-depleted cells, we hypothesize that Loqs regulates the efficiency of viral RNA replication.

The interaction of Loqs with DENV genomic RNA and viral protein NS3 suggests that Loqs can modulate the viral replication machinery. The enhanced binding of Loqs at the 3’ end of the viral negative strand may indicate a role for Loqs in the synthesis of positive viral RNA strands. However, the significance of the interaction of Loqs with sfRNAs is less clear. Alternatively, the relative affinity of Loqs for sfRNAs could be different than that for the genomic RNA, and these affinities may dictate distinct steps in the viral infectious cycle.

Loquacious is known to interact with R2D2, Ago2 and Dicer1/2 proteins to regulate siRNA and miRNA biogenesis [18]. While the mosquito midgut expresses isoforms that interact predominantly with Dicer to regulate siRNA generation and miRNA production, our experiments with Dicer-depleted cultured cells suggest that the pro-viral effects of Loqs is independent of any effects on small RNA biogenesis. A study showed that ectopic expression of Loqs2, a paralog of Loqs, inhibits systemic dissemination of DENV in in Aedes aegypti mosquitos by engaging the antiviral RNAi pathway [33]. The study also predicted a possible interaction between Loqs2 and Loqs in mediating this antiviral phenotype. In our study, we were unable to detect the endogenous Loqs2 mRNA in Aag2 cells. However, it is possible that Loqs is recruited by DENV to play a proviral role in tissues like the midgut where antiviral functions of the RNAi pathway are compromised. Future experiments that dissect the role of the RNAi pathway on viral replication in C3/36 and immuno-competent Aag2 cultured cells, and in different mosquito tissues will reveal the roles for canonical and non-canonical RNA binding proteins on flaviviral gene expression in the mosquito host.

## Materials and methods

### Cell culture

Huh7 and BHK21 cells were cultured as monolayers in Dulbecco’s modified Eagle’s medium. *Aedes albopictus* C6/36 cells and *Aedes aegypti* Aag2 cells were cultured as monolayers in Leibovitz’s L-15 and Schneider’s Drosophila media, respectively. All culture media were supplemented with 10% fetal bovine serum, 100 units of penicillin/ml, 100 µg of streptomycin/ml, 10 mM HEPES (pH 7.2), 1X NEAA (non-essential amino acid medium) and 2mM L-Glutamine (Gibco). Mammalian cell lines were grown at 37°C with 5% CO_2_, and insect cell lines were grown at 30°C without CO_2_. The Dicer2-knockout AF319 cell line was a kind gift from Dr. Kevin Maringer (The Pirbright Institute,UK).

### Plasmids

Plasmids pKF, pKF_EDEN15 and pKF_Scr containing BoxB stem loop sequences (GCCCTGAAAAAGGGC) flanking t3 repeats of the EDEN15 (TGTTTGTTTGTTTGTTGTTTGTTTGTTTGTTGTTTGTTTGTTTGT) or Scrambled EDEN15 (TTTTTGTTTTTTGGTTGTGGTTTTTTTTGTTGGGTTGGTTTTTTT) sequences and the BASU plasmid expressing the λN-biotin ligase fusion protein were a kind gift by Dr. Paul Khavari (Stanford University). A region of the DENV 3’UTR corresponding to sfRNA3 (271-455bp) was amplified from the DENV2-16681 cDNA and cloned in between BoxB sequences of pKF by Gibson assembly. The entire region containing GFP-BoxB-RNA of interest-BoxB-WPRE from pKF was subcloned into pBG34 (a kind gift from Dr. Brian Geiss, Colorado State University) by infusion cloning for expression in mosquito cells. The BG34-DENV 3’UTR clones used for RaPID experiments contain 2 BoxB sequences at the 5’ end and 3 BoxB sequences at the 3’ end of the DENV 3’UTR. The BG34-EDEN15 and BG34-Scr plasmids contain 3 flanking BoxB sequences both at their 5’ and 3’ ends. FLAG-tagged and HA-tagged pUB-Loqs and pUB-Loqs-PB plasmids were a kind gift by Dr. Zachary Adelman (Texas A&M University). EGFP was cloned into NdeI and SalI sites of the pUB vector to express HA-tagged GFP. The DENV2-New Guinea C strain and the DENV2-16681 luciferase reporter replicons were a kind gift by Dr. Jan Carette (Stanford University). Primers used in this study are listed in **Table S2**.

### In vitro RNA transcription

Dengue virus 2 serotype 16681 was propagated from infectious cDNA clone pD2IC/30P-A, a gift from Dr. Karla Kirkegaard (Stanford University). The DENV2 infectious cDNA and the replicon containing plasmids were linearized by digestion with XbaI and in vitro transcriptions were performed using the MEGAscript T7 transcription kit (AM1334). 5mg of the XbaI-linearized plasmid was incubated with 1.3 ml of 75mM rATP, 6.7 ml each of 75mM rCTP, rGTP and rUTP, 10 ml of 10X reaction buffer, 10 ml of T7 enzyme mix and 12.5 ml of 5’GpppA cap analog (S1406S, NEB) in a final reaction volume of 100 ml for 30 min at 30°C. 2.6 ml of 75mM rATP was added to the reaction and further incubated for 4 hrs at 30°C. The reactions were treated with DNAse, and RNA was purified using the RNEasy mini kit (Qiagen) according to manufacturer’s protocol.

### Virus generation and infection

In vitro transcribed capped DENV2 RNA was transfected into BHK21 cells in 24 well plates. Supernatants were collected 48 hrs post transfection and used to infect C6/36 cells overnight in a T75 flask containing 3ml complete medium. 15ml of complete medium was added to the flask the next day and virus supernatant was collected 6 days post infection. DENV2-NGC stocks were prepared similarly by infecting C6/36 cells in medium containing 2% FBS and HEPES. Viruses were titered on BHK21 cells to calculate PFU/ml. Virus infections were carried out by incubating cells with virus at the desired MOI for 1.5 hrs in 2% FBS-containing medium.

### Plaque assays

Dengue virus titers were measured using plaque assays on BHK-21 cells. Briefly, BHK-21 monolayers were grown to 90% confluence in 24-well plates and incubated with serially diluted virus supernatants for 1 hr at 37°C. The wells were subsequently overlaid with Dulbecco’s modified Eagle’s medium, 1% Aquacide and 5% FBS and incubated for 5-8 days. Cells were fixed with 10% formaldehyde for 20 min and stained with crystal violet for 20 min to visualize plaques. Plaque forming units (PFU) were calculated.

### Double stranded RNA preparation

Primers complementary to specific target gene sequences were designed using the E-RNAi website and the T7 promoter sequence was incorporated into both forward and reverse primers. The primers were used to amplify ∼500bp regions from target genes by PCR using cDNA extracted from Aag2 cells as template. PCR products were purified using the Qiagen PCR purification kit. In vitro transcription reactions (Promega MegaScript T7 transcription kit) containing ∼400 ng of the purified PCR product, 2 ml of 10X reaction buffer, 2 ml of each rNTP and 2 ml of T7 polymerase in a total volume of 20 ml were incubated overnight at 37°C. dsRNAs were DNAse-treated at 37°C for 30 min and purified using the RNEasy Mini kit. After annealing by heating at 95°C for 2min and slow cooling for 2hrs at 37°C aliquots were stored at −80°C.

### Transfection

For DENV2 virus generation, 1.5×10^5^ BHK-21 cells were seeded in each well of a 24-well plate and transfected with 1 mg of DENV2 RNA using Lipofectamine 3000. Medium was changed 2 hrs post transfection. For RaPID experiments, 10^7^ C6/36 cells were seeded onto a 10cm dish and co-transfected with 12 mg of pBG34-BoxB-RNA expressing plasmid and 1.5 mg of pBG34-BASU plasmid using 30 ml of Lipofectamine 3000. Transfection mix was prepared in 1ml OptiMEM and added to complete medium. Medium was changed 24 hrs post transfection. At 48 hrs post transfection, biotin was added to the medium at a final concentration of 10mM for 3 hrs. Cells were then harvested for RaPID experiments. For RNAi experiments, 2×10^5^ Aag2 cells were seeded in each well of a 24-well plate and transfected with 500 ng of dsRNA or 50 nM siRNA using 2 ml of Dharmafect-2 transfection reagent (Dharmacon). 100 ml of OptiMEM was used to prepare the transfection mix and added to 1 ml of complete medium in each well. Medium was changed at 6 hrs post transfection.

### RNA-protein interaction detection (RaPID)

10^7^ C6/36 cells were co-transfected with BoxB tagged-RNA and BASU expressing plasmids. 48 hrs post transfection, the medium was supplemented with biotin for 3 hrs. Cells were gently washed with cold 1X PBS on the plate, harvested and lysed with 600 μl lysis buffer (0.5M NaCl, 50mM Tris-HCl, 0.2% SDS, 1mM DTT) at room temperature. Next, 52 μl of 25% Triton X-100 was added to lysates and sonicated three times at an amplitude of 10% for 10 s at 30 s intervals. Further, 652 mL of cold 50mM Tris (pH 7.5) was added to lysates and briefly sonicated. Lysates were cleared by centrifugation at 14000 rpm for 10 min at 4°C. Clarified lysates were diluted in equal volume with 50mM Tris (pH 7.5) and centrifuged in 3k MWCO 15ml conical filters at 3900 rpm for 1 hr to remove free biotin. Supernatants from each filter were transferred into eppendorfs and protein concentrations were determined using Pierce Protein Quantitation Assay (ThermoFisher). Protein concentrations across samples were normalized to 4 mg/ml using 50mM Tris (pH 7.5). Biotinylated proteins were pulled down using MagResyn streptavidin beads with overnight rotation at 4°C (35 ml beads per mg protein). Beads were washed with a series of buffers for 5 min each at RT (Wash buffer 1: 2% SDS. Wash buffer 2: 0.1% Na-DOC, 1% Triton X-100, 0.5M NaCl, 50mM HEPES pH 7.5, 1mM EDTA. Wash buffer 3: 0.5% Na-DOC, 250μM LiCl, 0.5% NP-40, 10mM Tris-HCl, 1mM EDTA) and finally with 50mM Tris (pH 7.5). Washed beads were submitted to the Stanford Mass Spectrometry facility for downstream processing and LC-MS/MS analysis. Spectral counts of the identified peptides were filtered using CRAPome to eliminate background contaminants and probability scores were generated to identify peptides enriched in the experimental samples versus controls.

### qPCR

Total RNA was isolated from cells harvested in TRIzol (Invitrogen) according to manufacturer’s protocol. 1 mg of RNA was used for cDNA synthesis using High Capacity RNA-to-cDNA kit (Thermo Fisher, 4387406). 2 ml of cDNA was used to amplify target genes using the Power Up SyBR Green master mix (Thermo Fisher, A25742). Ct values of target genes were normalized to Ct values of the housekeeping gene, RPL32 to calculate fold-changes in RNA abundances.

### Luciferase assay

Cells were washed once with PBS and harvested in 100 ml of Renilla Luciferase Activity buffer (Promega). 10 μl aliquots were used to measure luminescence using the Luciferase Assay System (Promega) and the Glomax 20/20 luminometer with a 10 s integration time.

### Northern blot analysis

Total RNA was extracted from cells using TRIzol. 15μg RNA in RNA loading buffer (32% formamide, 1x MOPS-EDTA-Sodium acetate (MESA, Sigma) and 4.4% formaldehyde) was denatured at 65°C for 10 min and resolved on a 1% agarose gel containing 1x MESA and 3.7% formaldehyde. The RNA was transfered and UV crosslinked to a Zeta-probe membrane (Bio-Rad). Transfer efficiency was assessed by visualizing ribosomal RNA on the membrane using methylene blue stain. The membrane was destained and hybridized with α-^32^P dATP labelled DNA probes (RadPrime, Invitrogen) complementary to the DENV 3’UTR at 65°C for 3 hrs using ExpressHyb hybridization buffer (Clontech). Autoradiographs were quantified using ImageQuant (GE Healthcare).

### Western Blot

Cells were washed with PBS and lysed in RIPA buffer (50mM Tris (pH8.0),150 mM NaCl, 0.5% sodium deoxycholate, 0.1% SDS, and 1% Triton X-100) containing Complete EDTA-free protease inhibitors (Roche) for 30 min on ice. Cell lysates were clarified by centrifugation at 14000rpm for 5 min at 4°C. 50μg of cell lysate was mixed with 5x SDS loading dye (Thermo Fisher), denatured at 90°C for 10min and resolved on a 10% SDS-polyacrylamide gel. Proteins were transferred onto a PVDF membrane (Millipore), blocked with 5% non-fat milk in PBS-T and membranes were incubated with primary antibodies. Horse-radish peroxidase-conjugated secondary antibodies were used to visualize proteins using Pierce ECL Western Blot Substrate (Thermo Fisher) following manufacturer’s protocol. The following primary antibodies were used for western blot analysis: Anti-NS3 antibody (GTX124252, GeneTex), Anti-NS4B antibody (GTX124250, GeneTex), Anti-NS5 antibody (GTX103350, GeneTex), Anti-Capsid antibody (GTX103343, GeneTex), Anti-Actin antibody (A2066, Sigma), Anti-Loqs antibody (custom generated by GenScript), Anti-HA antibody (ab130275, Abcam) and Anti-GAPDH antibody (GTX627408, GeneTex).

### Polysome profile

10^7^ Aag2 cells were seeded onto 10cm plates and transfected with dsRNAs targeting GFP, Sec61A1 or Loqs genes. 24 hrs post transfection, cells were infected with DENV2-NGC at a MOI of 1. 48 h post infection, cells were treated for 3 min with cycloheximide (100 μg/mL) at 37°C, washed twice in cold PBS containing 100 μg/mL cycloheximide, and lysed for 10 min on ice in gradient buffer (150 mM KCl, 15 mM Tris–HCl, pH 7.5, 15 mM MgCl2, 100 μg/mL cycloheximide, 1 mg/mL heparin) containing 1% Triton X-100. Lysates were cleared by centrifugation at 14000rpm for 10 min at 4°C and layered onto 10% to 60% sucrose gradients composed of the above gradient buffer. Gradients were spun in an SW41 ultra-centrifuge rotor for 2 h 45 min at 35,000 rpm at 4°C. Fractions were collected using the Isco Retriever II/UA-6 detector system. RNA was isolated from each fraction using the RNEasy mini-kit and used for cDNA preparation and qPCR analysis.

### RNP immunoprecipitation

10^7^ Aag2 cells were seeded onto 10cm plates and transfected with plasmids expressing HA-GFP or HA-Loqs PA/PB. 24 hrs post transfection, cells were infected with DENV2-NGC at a MOI of 1. 48 hrs post infection, cells were washed with cold 1X PBS on the plate and harvested in 1 ml of Pierce IP lysis buffer containing protease inhibitors. Lysates were incubated on ice for 30 min, clarified by centrifugation at 14000rpm for 10 min at 4°C and protein concentrations in the samples were estimated by Bradford assay. For each sample, a total of 400 mg protein at a concentration of <1 mg/ml was precleared by rotating with 10 m l of Protein G Dynabeads for 1 hr at 4°C. Lysates were subsequently incubated with Anti-HA Dynabeads overnight at 4°C with rotation. RNase treatment was performed by incubating lysates with RnaseA/T1 (0.4 U/ml RNase A and 16.6 U/ml RNase T1) for 15 min at 37°C prior to addition of anti-HA beads. For immunoprecipitating endogenous Loqs, lysates were incubated with beads saturated overnight with 4 mg of anti-Loqs antibody or a rabbit IgG isotype control. The following day, beads were washed thrice with cold Pierce IP lysis buffer and twice with the same buffer supplemented with 500mM NaCl. For RNA elution, beads were treated with 30 mg of ProteinaseK in IP lysis buffer containing 0.1% SDS. RNA was extracted using TRIzol-LS and analyzed by northern blot or RT-qPCR. For protein co-immunoprecipitation experiments, proteins were eluted from beads by boiling with 1X SDS gel loading buffer and analyzed by Western blot.

### Detergent fractionation of cells

2×10^6^ Aag2 cells were seeded onto each well of a 6 well plate and infected with DENV at a MOI of 1 for 48 hrs. Cells were gently washed on the plate with 3ml of cold PBS, harvested in 1ml PBS and pelleted by centrifugation at 1000g for 5 min at 4°C. Next, cells were lysed by resuspending in 1 ml permeabilization buffer (110mM KOAc, 25mM HEPES-KOH (pH-7.5), 2.5mM Mg(OAc)2, 1mM EGTA, 0.015% digitonin, 1mM DTT, 40U/ml RNAseOUT), incubated for 5 min at 4°C and pelleted as above. Supernatants (cytosolic fraction) were transferred into fresh Eppendorf tubes and remaining pellets were washed by resuspension in 5 ml of wash buffer (110mM KOAc, 25mM HEPES-KOH (pH 7.5), 2.5mM Mg(OAc)2, 1mM EGTA, 0.004% digitonin, 1mM DTT) and pelleted as above. Washed pellets were resuspended in 250 ml lysis buffer (400mM KOAc, 25mM HEPES-KOH pH 7.5, 15mM Mg(OAc)2, 1% (v/v) NP-40, 0.5% (w/v) sodium deoxycholate, 1mM DTT) to solubilize the membrane fractions, incubated for 5min at 4°C and centrifuged at 7500g for 10 min at 4°C. Supernatants (membrane fraction) were transferred into fresh tubes while the remaining pellets were saved (nuclear fraction). 20 ml from each fraction was analyzed by western blot.

### RNA fluorescence in situ hybridization (RNA-FISH)

RNA-FISH was performed using the RNA View Cell Plus assay kit (Cat. No.88-19000-99, ThermoFisher) according to the manufacturer’s protocol. Briefly, 2×10^5^ Aag2 cells were plated onto coverslips in 24 well plates, infected with DENV at a MOI of 1 and fixed using 4% paraformaldehyde for 30 min at RT. Cells were permeabilized, blocked and incubated with Anti-Loqs or anti-NS3 primary antibodies at a dilution of 1:200 followed by incubation with Alexa-Fluor 488 secondary antibodies (Invitrogen) at a dilution of 1:500. After antibody staining, cells were washed with PBS and incubated with DNA probes complementary to the NS5 region of DENV genomic RNA at 40°C for 2 hrs (1:100 dilution). Cells were then sequentially treated at 40°C for 1 hr with the pre-amplifier mix, amplifier mix and Label Probe mix and finally mounted on slides using Fluoromount-G with DAPI. Imaging analysis was carried out at the Stanford Imaging Facility.

### Infrared UV-crosslinking immunoprecipitation (irCLIP) of Loqs

10^7^ Aag2 cells were seeded onto 10cm plates, transfected the next day with HA-GFP or HA-Loqs PA/PB plasmids and infected the following day with DENV2-NGC at MOI of 1. After 48 hrs, infected cells were UV crosslinked at 0.35 J/cm2, lysed in CLIP lysis buffer (50 mM HEPES, 200 mM NaCl, 1 mM EDTA, 10% glycerol, 0.1% NP-40, 0.2% Triton X-100, 0.5% N-lauroylsarcosine). Isolation and processing of RNA-protein complexes were performed as described [13]. Briefly, sequential immunoprecipitations were performed using the anti-HA antibody followed by the anti-Loqs antibody for 8 hours at 4°C on rotation. RNP-complexes were resolved on SDS-PAGE gels, transferred onto nitrocellulose, excised and the RNA isolated for library preparation. The Dengue genome was downloaded from NCBI (GenBank: AF038403.1) and the mosquito genome (AaegL5) was downloaded from Ensembl. Genomes were indexed using hisat2-build. Trimmed reads were mapped to the dengue genome and then the mosquito genome using hisat2. Bam files from the three replicates were merged and then visualized using the Integrative Genomics Viewer. For the dengue genome mapping results, reads mapping to the forward and reverse strand were separated using “samtools view -F 20” and “samtools view -f 16” and then viewed in IGV.

## Acknowledgements

We are grateful to Karla Kirkegaard for many comments and critical reading of the manuscript. We thank C. Adams, R. Leib and the Vincent Coates Foundation Mass Spectrometry Laboratory, Stanford University Mass Spectrometry for help with mass spectrometry.

## Author Contributions

**Conceptualization:** Peter Sarnow, Shwetha Shivaprasad

**Funding acquisition:** Peter Sarnow, Ryan Flynn

**Formal analysis:** Shwetha Shivaprasad, Kuo-Feng Weng, Ryan Flynn, Julia Belk, Peter Sarnow

**Investigation:** Shwetha Shivaprasad, Kuo-Feng Weng, Ryan Flynn, Julia Belk, Peter Sarnow

**Project administration:** Peter Sarnow, Ryan Flynn

**Supervision:** Peter Sarnow, Ryan Flynn

**Writing – original draft:** Shwetha Shivaprasad

**Writing -review & editing:** Shwetha Shivaprasad, Kuo-Feng Weng, Ryan Flynn, Julia Belk, Peter Sarnow

## Supplementary Figures

**Fig S1.**
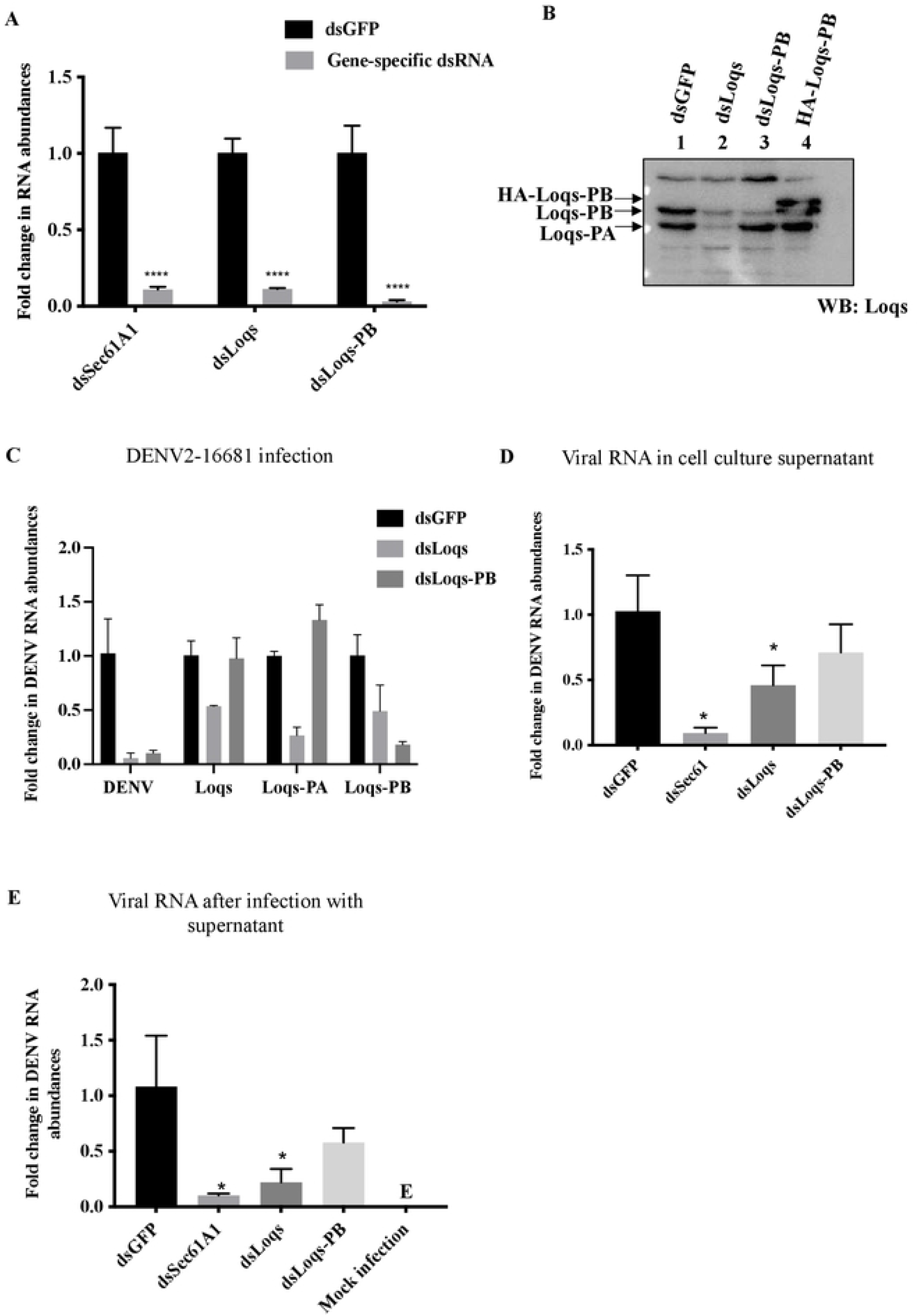
Effects of Loqs isoform depletion on DENV2-16681 (Thailand strain). (A) RT-qPCR measurement of mRNA abundances in Aag2 cells transfected with the indicated dsRNAs. Knockdown efficiency was measured using gene-specific primers (n=3, **p<0.005). (B) Western blot analysis of Loqs protein abundance in dsGFP, dsLoqs, dsLoqs-PB and HA-Loqs PB transfected Aag2 cells (C) Effects of dsRNA treatment on DENV2-16681(Thailand strain) infection of Aag2 cells (MOI=0.1, 96 hrs), measured by RT-qPCR. Knockdown efficiency was measured using gene-specific primers. Measurements are represented as fold-change over dsGFP (n=3, *p<0.05, **p<0.005). (D) Effect of dsRNA treatment on extracellular abundances of DENV2-16681 viral RNA in infected Aag2 cells, measured by RT-qPCR. Data is plotted as fold-change over dsGFP from three independent experiments. (E) Cell culture supernatants from dsRNA-treated cells were used to infect naive Aag2 cells and viral RNA abundances in these cells were measured by RT-qPCR.

**Fig S2.**
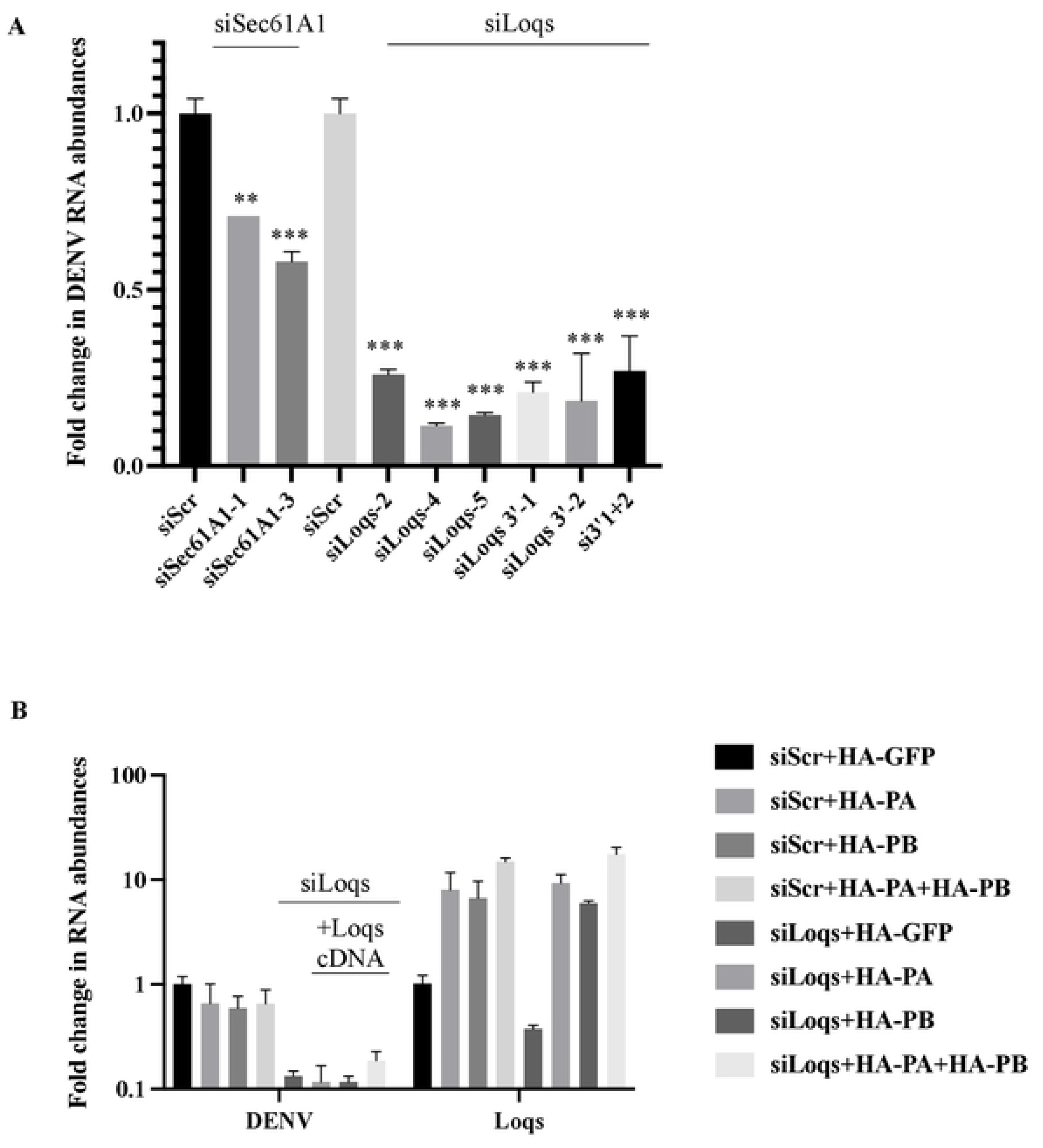
Effects of over-expression of different Loqs isoforms and different Loqs siRNAs on DENV2 RNA abundances. (A) Aag2 cells were transfected with the indicated siRNAs for 24 hrs followed by DENV2 infection. Cells were harvested for qPCR at 96 hrs post infection. Intracellular DENV2 RNA abundances are represented as average fold-change over siScr from three independent experiments. (B) Aag2 cells were co-transfected with scrambled (siScr) or Loqs (siLoqs 3’-2) siRNAs and the indicated plasmid DNAs. 24 hrs post transfection they were infected with DENV2-NGC virus at a MOI of 0.1. Cells were harvested at 96 hrs post infection. Intracellular DENV2 RNA abundances were measured by RT-qPCR and are represented as average fold-change over the siScr from three independent experiments.

**Fig S3.**
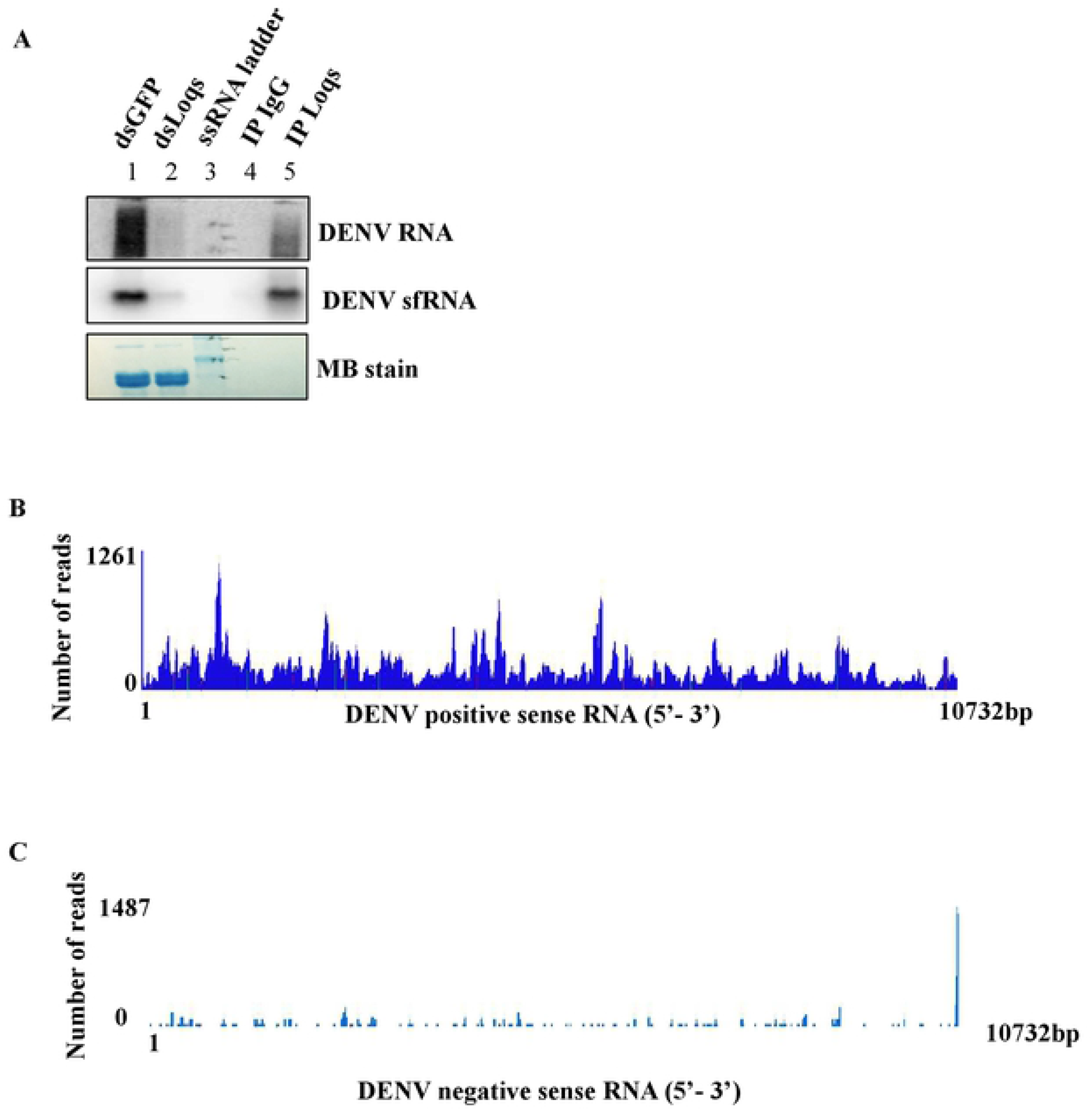
Infrared UV-crosslinking immunoprecipitation (irCLIP) of Loqs. (A) Northern blot analysis of RNA immunoprecipitated from DENV (MOI=1) infected Aag2 cell lysates using Loqs antibody or a control antibody. DENV genomic RNA and DENV subgenomic RNAs (DENV-sfRNA) were detected using radiolabeled RNA probes. Methylene blue (MB) staining of rRNA is shown as a loading control. Representative image from three independent immunoprecipitation experiments is shown. (B,C) Aag2 cells were transfected with HA-GFP or HA-Loqs PA/PB plasmids. 24 hrs post transfection, cells were infected with DENV2-NGC at a MOI of 1. Cells were UV irradiated at 254nm to covalently crosslink RNA-protein interactions and subjected to irCLIP with anti-Loqs followed by anti-HA antibodies. irCLIP RT stops were mapped at base resolution to the DENV genome. The read density across positive-(B) and negative-(C) sense DENV RNAs is represented as an average of three independent experiments.

**Fig S4.**
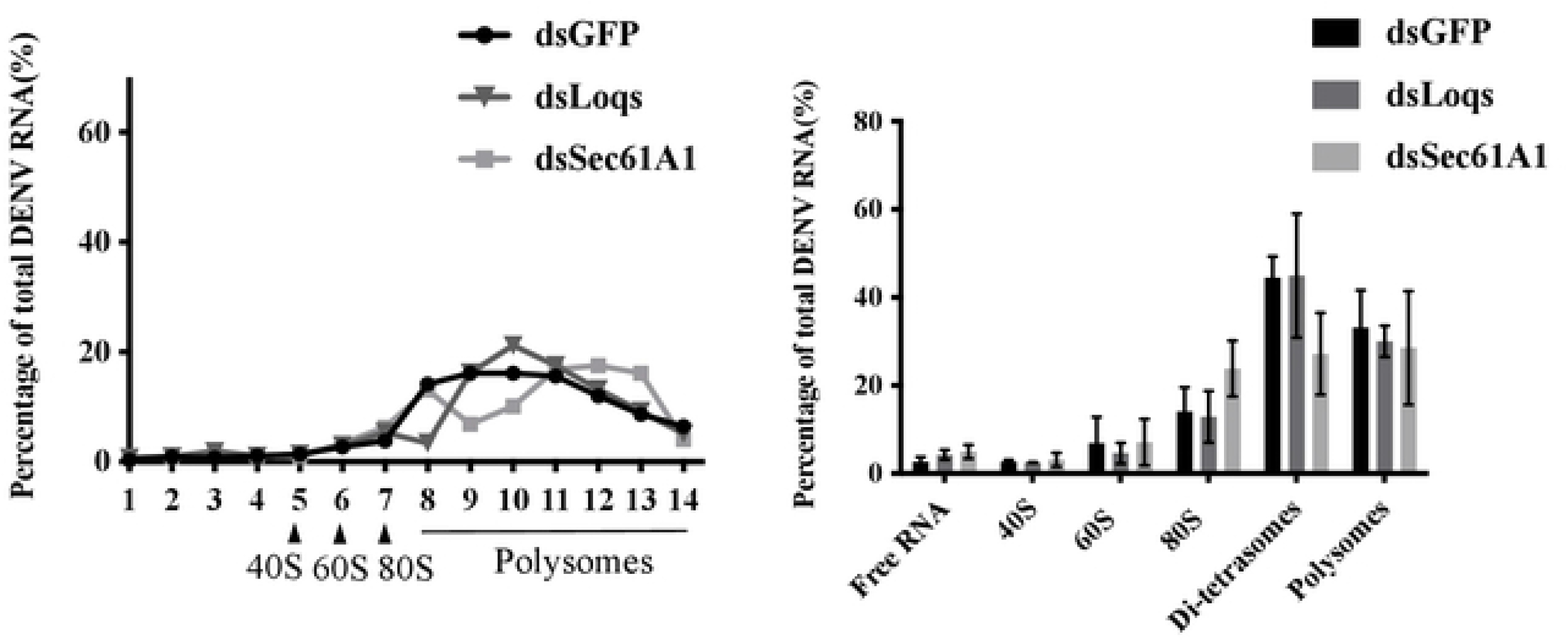
Polysome analysis of dsRNA-treated Aag2 cells infected with DENV2 at an MOI of 1 for 48h. (A) DENV2 RNA abundance in each polysome fraction was measured by RT-qPCR and plotted as a percentage of the total RNA. A representative graph from three independent experiments is shown. (B) DENV2 RNA abundance in indicated polysome fractions plotted as an average from three independent experiments.

**Fig S5.**
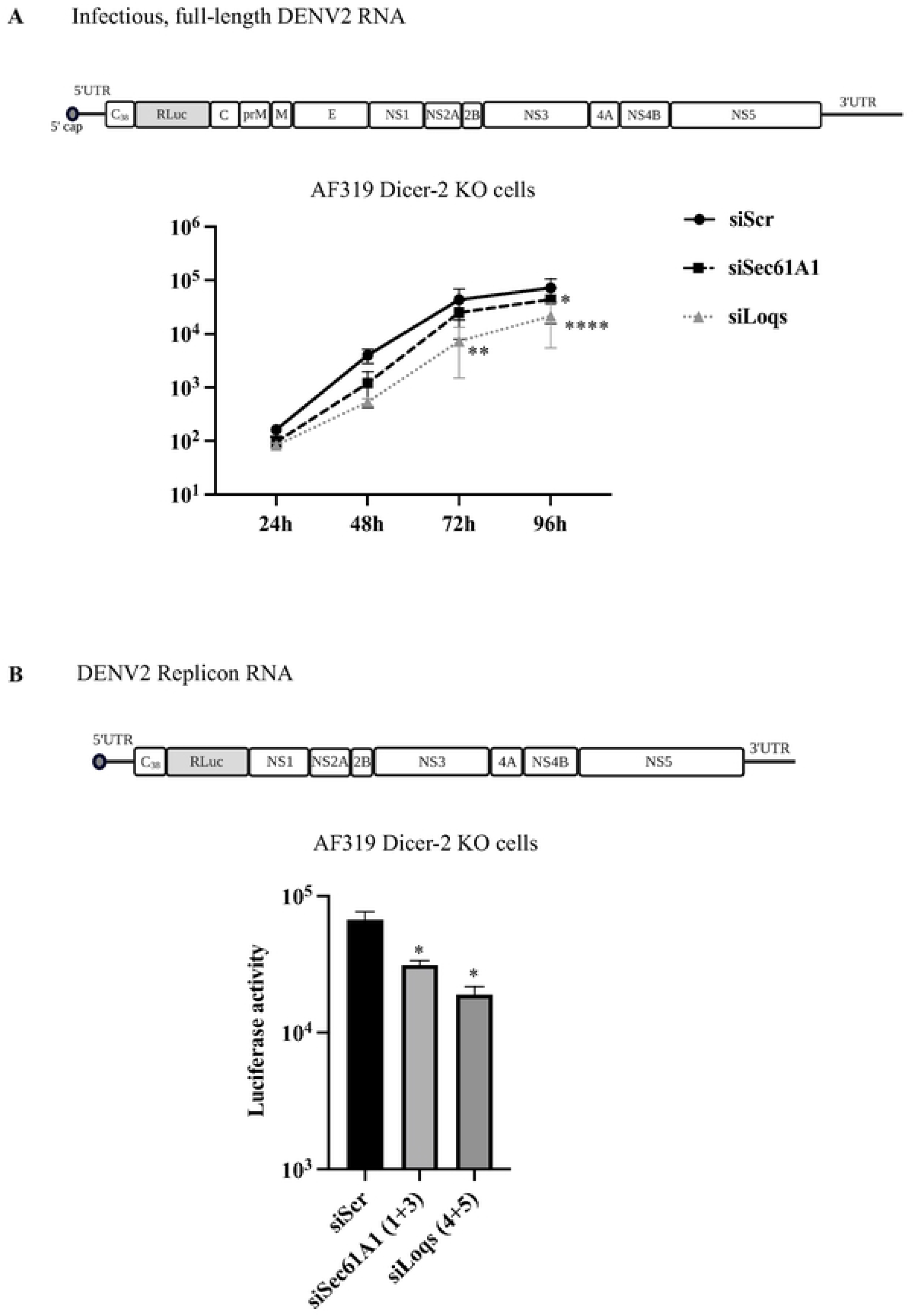
Effect of Dicer depletion on Loqs inhibition of DENV replication. (A) AF319 Dicer-2 knock-out (KO) cells were transfected with the indicated siRNAs. 24 hrs after siRNA transfection, cells were infected with luciferase expressing DENV2 virus. Luciferase expression in cell lysates was measured at the indicated time points and represented as an average from three independent experiments. (B) AF319 Dicer-2 KO cells were transfected with the indicated siRNAs. At 24 hrs after siRNA transfection, cells were transfected with luciferase expressing DENV2-NGC replicon RNAs. Luciferase expression in cell lysates was measured 96 hrs post transfection.

